# Leg length, skull circumference, and the incidence of dementia in Latin America and China; a 10/66 population-based cohort study

**DOI:** 10.1101/140129

**Authors:** Martin J. Prince, Daisy Acosta, Mariella Guerra, Yueqin Huang, Ivonne Z Jimenez-Velazquez, Juan J. Llibre Rodriguez, Aquiles Salas, Ana Luisa Sosa, Michael E. Dewey, Maelenn M. Guerchet, Zhaorui Liu, Jorge J. Llibre-Guerra, Matthew A. Prina

## Abstract

**Background:** Adult leg length is influenced by nutrition in the first few years of life. Adult head circumference is an indicator of brain growth. Cross-sectional studies indicate inverse associations with dementia risk, but there have been few prospective studies.

**Methods:** Population-based cohort studies in urban sites in Cuba, Dominican Republic Puerto Rico and Venezuela, and rural and urban sites in Peru, Mexico and China. Sociodemographic and risk factor questionnaires were administered to all participants, and anthropometric measures taken, with ascertainment of incident 10/66 dementia, and mortality, three to five years later.

**Results:** Of the original at risk cohort of 13,587 persons aged 65 years and over, 2,443 (18.0%) were lost to follow-up; 10,540 persons with skull circumference assessments were followed up for 40,466 person years, and 10,400 with leg length assessments were followed up for 39,954 person years. There were 1,009 cases of incident dementia, and 1,605 dementia free deaths. The fixed effect pooled meta-analysed adjusted subhazard ratio (ASHR) for leg length (highest vs. lowest quarter) was 0.80 (95% CI, 0.66-0.97) and for skull circumference was 1.02 (95% CI, 0.84-1.25), with no heterogeneity of effect between sites (I^2^=0%). Leg length measurements tended to be shorter at follow-up, particularly for those with baseline cognitive impairment and dementia. However, leg length change was not associated with dementia incidence (ASHR, per cm 1.006, 95% CI 0.992-1.020), and the effect of leg length was little altered after adjusting for baseline frailty (ASHT 0.82, 95% CI 0.67-0.99). A priori hypotheses regarding effect modification by gender or educational level were not supported.

**Conclusions:** Consistent findings across settings provide quite strong support for an association between adult leg length and dementia incidence in late-life. Leg length is a relatively stable marker of early life nutritional programming, which may confer brain reserve and protect against neurodegeneration in later life through mitigation of cardiometabolic risk. Further clarification of these associations could inform predictive models for future dementia incidence in the context of secular trends in adult height, and invigorate global efforts to improve childhood nutrition, growth and development.

## Introduction

The foetal and developmental origins of adult disease may also be relevant to the aetiology of dementia [1,2]. Late-life leg length and skull circumference provide useful information about the nutritional environment in early life, and brain development, respectively. Adult leg length is sensitive to diet in infancy, specifically breast feeding and energy intake [3]. Increasing population height is mainly accounted for by increasing leg length, linked to improvements in childhood nutrition [4]. Skull dimensions are a long-term and stable marker of early-life brain size [5,6].

Inverse associations between skull circumference and prevalent Alzheimer’s disease (AD) were reported in five cross-sectional studies: three of which were community based, from the USA [7], Brazil [8], and Korea [9], and two of communities of Catholic nuns from the US [10] and Germany [11]. There are two negative reports [12,13], both small case-control studies with cases recruited from clinical services. In the only cohort study an inverse association was noted with incident AD among Japanese-Americans [14]. Leg length was inversely cross-sectionally related to dementia prevalence in Brazil [8], and among women in a population-based study in Korea [15], and with cognitive impairment among Caribbean migrants to the UK [16]. In the only cohort study, from the USA, knee height was inversely associated with incident dementia, but among women only [17]. This was a post hoc finding, and the interaction with gender was not statistically significant.

In the baseline phase of the 10/66 Dementia Research Group studies in Latin America, China and India, longer legs and larger skulls were independently associated with a lower prevalence of dementia [18]. There was little heterogeneity of estimates among sites. Earlier reports of effect modification were not replicated [9,10,17]. The incidence phase of the 10/66 Dementia Research Group surveys allows us to

1. test hypotheses that longer leg lengths and larger skull circumferences are associated with a lower incidence of dementia in large, representative population-based dementia-free cohorts, checking for consistency across diverse urban and rural settings in Latin America, and China
2. test for effect modification, with a priori hypotheses based upon previously observed associations

a. any protective effect of longer leg length on dementia incidence is modified by gender [9,15,17] (stronger in women)
b. any protective effect of greater skull circumference on dementia incidence is modified by education [10] (stronger among the least educated), or gender [9] (stronger in women)
3. assess the possibility that reverse causality may have accounted for previously observed cross-sectional associations by testing whether

a. cognition at baseline is associated with changes in leg and skull measurements from baseline to follow-up
b. reduction in leg lengths and skull circumferences are associated with the incidence of dementia.

## Materials and Methods

The 10/66 population-based baseline and incidence wave study protocols are published in an open access journal [19], and the profile of the resultant cohort has also been described [20]. Relevant details are provided here. One-phase population-based surveys were carried out of all residents aged 65 years and over in geographically defined catchment areas (urban sites in Cuba, Dominican Republic, Puerto Rico and Venezuela, and urban and rural sites in Mexico, Peru, and China)[19]. Baseline surveys were completed between 2003 and 2007, other than in Puerto Rico (2007-2009). The target sample was 2000 for each country, and 3000 for Cuba. The baseline survey included clinical and informant interviews, and physical examination. Incidence waves were subsequently completed, with a mortality screen, between 2007 and 2011 (2011-2013 in Puerto Rico) aiming for 3-4 years follow-up in each site [21]. Assessments were identical to baseline protocols for dementia ascertainment, and similar in other respects. We revisited participants’ residences on up to five occasions. If no longer resident, we sought information on their vital status and current residence from contacts recorded at baseline. If moved away, we sought to re-interview them, even outside the catchment area. If deceased, we recorded the date, and completed an informant verbal autopsy, including evidence of cognitive and functional decline suggestive of dementia onset between baseline assessment and death [22].

## Measures

The 10/66 survey interview generates information regarding dementia diagnosis, mental disorders, physical health, anthropometry, demographics, an extensive dementia and chronic diseases risk factor questionnaire, disability, and health service utilisation [19]. Only assessments relevant to the analyses of associations between leg length, skull circumference and dementia incidence are described in detail here.

### Anthropometric measures

Skull circumference was measured using a cloth tape measure encircling the nuchal tuberosity and the brow. Leg length was measured, standing, from the highest point of the iliac crest to the lateral malleolus. Both dimensions were measured, to the nearest centimeter, at both time points.

### Sociodemographic variables

Age was determined at baseline interview from stated age, documentation and informant report, and, if discrepant, according to an event calendar. We also recorded sex, and education level (none/ did not complete primary/ completed primary/ secondary/ tertiary).

### Frailty

At baseline, we assessed four of the five indicators of the Fried physical frailty phenotype [23]; exhaustion, weight loss, slow walking speed and low energy expenditure. Since hand grip strength was not measured we considered participants as frail if they fulfilled two or more of the four frailty indicators [24].

### Dementia

10/66 dementia diagnosis is allocated to those scoring above a cutpoint of predicted probability for dementia, calculated using coefficients from a logistic regression equation developed, calibrated and validated cross-culturally in the 25 centre 10/66 pilot study[25], applied to outputs from a) a structured clinical interview, the Geriatric Mental State[26], b) two cognitive tests; the Community Screening Instrument for Dementia (CSI-D) COGSCORE[27] and the modified CERAD 10 word list learning task delayed recall[28], and c) informant reports of cognitive and functional decline from the CSI-D RELSCORE[27]. For those who died between baseline and follow-up we diagnosed ‘probable incident dementia’ by applying three criteria:

1. A score of more than two points on the RELSCORE, from the post-mortem informant interview, with endorsement of either ‘deterioration in memory’ or ‘a general deterioration in mental functioning’ or both, and
2. an increase in RELSCORE of more than two points from baseline, and
3. the onset of these signs noted more than six months prior to death.

In the baseline survey, the first criterion would have detected those with either DSM-IV or 10/66 dementia with 73% sensitivity and 92% specificity[22].

Prevalence[29] and incidence[22] of 10/66 dementia in the current cohorts has been reported.

### Analyses

We used STATA version 11 for all analyses.

For each site

1. we describe participants’ characteristics; age, sex, educational level, mean leg length, and skull circumference
2. we model the effect of covariates on 10/66 dementia incidence using a competing-risks regression derived from Fine and Gray’s proportional subhazards model [30]. This is based on a cumulative incidence function, indicating the probability of failure (dementia onset) before a given time, acknowledging the possibility of a competing event (case dementia-free death). In our analysis of baseline data, men had larger skulls and longer legs than women in all sites, while in most sites older people had smaller skulls and shorter legs, consistent with birth cohort effects [18]. Better education was generally associated with larger skulls and longer legs. We therefore controlled for age, gender, and education when modelling the effects of leg length (quarters) and skull circumference (quarters) on 10/66 dementia incidence, comparing the longest or largest quarters with the shortest or smallest. We report fully adjusted sub-hazard ratios (ASHR) with robust 95% confidence intervals adjusted for household clustering. As a sensitivity analysis, we also estimate the linear effect of leg length and skull circumference (per centimetre) on dementia incidence, controlling for the same covariates. We fit the models separately for each site and then use a fixed effects meta-analysis to combine them. Higgins I^2^ quantifies the proportion of between-site variability accounted for by heterogeneity, as opposed to sampling error; up to 40% heterogeneity is conventionally considered negligible, while up to 60% may reflect moderate heterogeneity [31].
3. we test for effect modification of linear effects of leg length on 10/66 dementia by gender, and of skull circumference upon 10/66 dementia by gender and education, by extending the adjusted models by appropriate interaction terms.
4. we estimate correlations between baseline and follow-up measures of skull circumference and leg length, and explore the possibility of reverse causation by a) comparing change in leg length and skull circumference by baseline cognitive status (cognitively normal, versus ‘cognitive impairment no dementia (CIND), versus dementia), and b) testing whether change in leg length or skull circumference is associated with incident dementia.

Participation in the study was by signed informed consent, or by signed independently witnessed verbal consent for participants who could not read or write. The study protocol and the consent procedures were approved by the King's College London research ethics committee and in all countries where the research was carried out: 1- Medical Ethics Committee of Peking University the Sixth Hospital (Institute of Mental Health, China); 2- the Memory Institute and Related Disorders (IMEDER) Ethics Committee (Peru); 3- Finlay Albarran Medical Faculty of Havana Medical University Ethical Committee (Cuba); 4- Hospital Universitario de Caracas Ethics Committee (Venezuela); 5- Consejo Nacional de Bioética y Salud (CONABIOS, Dominican Republic); 6- Instituto Nacional de Neurología y Neurocirugía Ethics Committee (Mexico); 7- University of Puerto Rico, Medical Sciences Campus Institutional Review Board (IRB)

## Results

### Sample characteristics

Across the 10 sites, 15,027 interviews were completed at baseline. Response proportions varied between 72% and 98% [29]. The ‘at risk’ cohort comprised 13,587 dementia-free persons aged 65 years and over (Table 1). Mean age at baseline varied from 72.0 to 75.4 years, lower in rural than urban sites and in China than in Latin America. Women predominated over men in all sites, accounting for between 53% and 67% of participants, by site. Educational levels were lowest in rural Mexico (83% not completing primary education), Dominican Republic (70%), rural China (69%), and urban Mexico (54%) and highest in urban Peru (9%), Puerto Rico (20%), and Cuba (23%). In other sites, between a third and a half of participants had not completed primary education. There was significant between site variation (Table 1) in leg length (shorter legs in rural compared with urban sites; F=64.7, p<0.04), and skull circumference (no obvious pattern of variation; F=82.3, p<0.001). Site accounted for 5.5% of the variance in skull circumference, and 4.4% of the variance in leg length. From the ‘at risk’ cohort, 2,443 participants (18.0%) were lost to follow-up; 10,540 with baseline skull circumference data were followed up for 40,466 person years, and 10,400 with baseline leg length data were followed up for 39,954 person years (Table 1). In the skull circumference cohort, there were 1,766 deaths, of which 1,605 were judged to be dementia-free, and 1,009 cases of incident dementia were observed (of which, 161 were ‘probable’ cases diagnosed retrospectively from post-mortem informant interviews).

**Table 1.** Cohort characteristics, by site

| Site | At risk (n) | Status at follow up |  | Skull circumference |  | Leg length |  | Outcome (skull circumference analysis) |  |
| --- | --- | --- | --- | --- | --- | --- | --- | --- | --- |
|  |  | Status | Number (%) | Mean (SD) | Numbers in cohort analysis (pyears) | Mean (SD) | Numbers in cohort analysis (pyears) | Outcome | Number (%) |
| Cuba | 2621 | Interviewed | 1851 (70.6%) | 55.9 (1.9)<br>MV=41 | 2259 (9134) | 85.3 (7.4)<br>MV=95 | 2205 (8900) | Censored | 1649 (73.0%) |
|  |  | Deceased | 449 (17.1%) |  |  |  |  | Incident dementia | 181 (8.0%) |
|  |  | Lost | 321 (12.2%) |  |  |  |  | Competing risk | 429 (19.0%) |
| Dominican Republic | 1769 | Interviewed | 1071 (60.6%) | 56.3 (2.3)<br>MV=31 | 1410 (6068) | 87.3 (6.8)<br>MV=53 | 1388 (5968) | Censored | 936 (66.4%) |
|  |  | Deceased | 370 (20.9%) |  |  |  |  | Incident dementia | 162 (11.5%) |
|  |  | Lost | 328 (18.5%) |  |  |  |  | Competing risk | 312 (22.1%) |
| Puerto Rico |  | Interviewed | 1174 (66.5%) | 54.9 (2.1)<br>MV=188 | 1186 (4849) | 89.4 (7.1)<br>MV=203 | 1171 (4781) | Censored | 926 (78.1%) |
|  |  | Deceased | 200 (11.3%) |  |  |  |  | Incident dementia | 131 (11.0%) |
|  |  | Lost | 391 (22.2%) |  |  |  |  | Competing risk | 129 (10.9%) |
| Peru Urban | 1251 | Interviewed | 819 (65.5%) | 55.3 (2.1)<br>MV=17 | 863 (2431) | 87.6 (7.8)<br>MV=47 | 833 (2361) | Censored | 762 (88.3%) |
|  |  | Deceased | 61 (4.9%) |  |  |  |  | Incident dementia | 43 (5.0%) |
|  |  | Lost | 371 (29.7%) |  |  |  |  | Competing risk | 58 (6.7%) |
| Peru Rural | 516 | Interviewed | 395 (76.5%) | 55.8 (2.1)<br>MV=9 | 434 (1489) | 84.2 (7.2)<br>MV=14 | 429 (1471) | Censored | 362 (83.4%) |
|  |  | Deceased | 48 (9.3%) |  |  |  |  | Incident dementia | 32 (7.4%) |
|  |  | Lost | 73 (14.1%) |  |  |  |  | Competing risk | 40 (9.2%) |
| Venezuela | 1820 | Interviewed | 1192 (65.5%) | 55.2 (2.4)<br>MV=270 | 1083 (4318) | 86.9 (9.2)<br>MV=281 | 1072 (4294) | Censored | 847 (78.2%) |
|  |  | Deceased | 161 (8.8%) |  |  |  |  | Incident dementia | 127 (11.7%) |
|  |  | Lost | 467 (25.7%) |  |  |  |  | Competing risk | 109 (10.1%) |
| Mexico Urban | 910 | Interviewed | 700 (76.9%) | 55.0 (2.1)<br>MV=15 | 763 (2182) | 85.8 (6.7)<br>MV=11 | 767 (2197) | Censored | 644 (84.4%) |
|  |  | Deceased | 78 (8.6%) |  |  |  |  | Incident dementia | 45 (5.9%) |
|  |  | Lost | 132 (14.5%) |  |  |  |  | Competing risk | 74 (9.7%) |
| Mexico Rural | 913 | Interviewed | 655 (71.8%) | 54.4 (1.9)<br>MV=6 | 737 (2182) | 82.7 (6.2)<br>MV=13 | 730 (1997) | Censored | 573 (77.7%) |
|  |  | Deceased | 88 (9.6%) |  |  |  |  | Incident dementia | 83 (11.3%) |
|  |  | Lost | 170 (18.6%) |  |  |  |  | Competing risk | 81 (11.0%) |
| China Urban | 1076 | Interviewed | 711 (66.1%) | 55.2 (2.0) | 877 (3914) | 88.1 (7.2) | 864 (3859) | Censored | 622 (70.9%) |
|  |  | Deceased | 175 (16.3%) |  |  |  |  | Incident dementia | 109 (12.4%) |
|  |  | Lost | 190 (17.7%) | MV=9 |  | MV=22 |  | Competing risk | 146 (16.6%) |
| China Rural | 946 | Interviewed | 695 (73.4%) | 55.0<br>(2.0)<br>MV=18 | 928<br>(4066) | 86.9<br>(11.4)<br>MV=5 | 941<br>(4126) | Censored | 605 (65.2%) |
|  |  | Deceased | 251 (26.5%) |  |  |  |  | Incident dementia | 96 (10.3%) |
|  |  | Lost | 0 (0.0%) |  |  |  |  | Competing risk | 227 (24.5%) |
| Total | 13587 | Interviewed | 9263 (68.2%) | 55.4<br>(2.4)<br>MV=604 | 10540<br>(40466) | 86.6<br>(8.0)<br>MV=744 | 10400<br>(39954) | Censored | 7926 (75.2%) |
|  |  | Deceased | 1881 (13.8%) |  |  |  |  | Incident dementia | 1009 (9.6%) |
|  |  | Lost | 2443 (18.0%) |  |  |  |  | Competing risk | 1605 (15.2%) |

### Leg length and incident dementia

In all sites other than Cuba, there was a non-significant trend towards inverse associations between leg length and incident dementia (Table 2). The pooled meta-analysed adjusted sub-hazard ratio (ASHR) for leg length (longest vs. shortest quarter) was 0.80 (95% CI, 0.66-0.97) with no heterogeneity between sites (I^2^=0%). The linear effect, per centimetre of leg length, was also statistically significant when pooled across sites (SHR 0.986, 95% CI 0.977-0.995, I^2^=0%). The interaction between leg length (per quarter) and gender (male versus female) was not statistically significant (pooled ASHR 1.03, 95% CI 0.90-1.18, I^2^=38.3%).

**Table 2.** Associations of skull circumference and leg length with incident 10/66 dementia (competing risk^1^ proportional hazards regression)

| Site | Skull circumference |  | Leg length |  |
| --- | --- | --- | --- | --- |
|  | Adjusted<sup>2</sup> SHR<sup>3</sup><br>Largest quarter<br>vs smallest<br>quarter | Adjusted<sup>2</sup> SHR<br>Linear effect (per<br>cm) | Adjusted<sup>2</sup> SHR<br>Longest quarter<br>vs shortest<br>quarter | Adjusted<sup>2</sup> SHR<br>Linear effect (per<br>cm) |
| Cuba | 1.69 (0.93-3.07) | 1.033 (0.956-1.115) | 1.17 (0.74-1.87) | 1.005 (0.986-1.026) |
| Dominican<br>Republic | 0.87 (0.53-1.43) | 0.992 (0.929-1.058) | 0.65 (0.40-1.08) | 0.969 (0.947-0.993) |
| Puerto Rico | 0.99 (0.57-1.72) | 0.988 (0.907-1.077) | 0.90 (0.51-1.58) | 0.992 (0.953-1.031) |
| Peru Urban | 1.34 (0.50-3.58) | 1.091 (0.910-1.309) | 0.89 (0.38-2.10) | 0.993 (0.952-1.036) |
| Peru Rural | 1.12 (0.40-3.11) | 0.916 (0.734-1.145) | 0.45 (0.10-2.01) | 0.962 (0.922-1.003) |
| Venezuela | 0.93 (0.55-1.55) | 0.992 (0.922-1.067) | 0.75 (0.43-1.30) | 0.980 (0.956-1.005) |
| Mexico Urban | 1.48 (0.58-3.73) | 0.890 (0.766-1.034) | 0.72 (0.32-1.63) | 0.975 (0.934-1.019) |
| Mexico Rural | 0.96 (0.54-1.71) | 1.002 (0.898-1.118) | 0.86 (0.46-1.62) | 0.978 (0.938-1.021) |
| China Urban | 1.43 (0.74-2.79) | 1.040 (0.923-1.172) | 0.66 (0.38-1.16) | 0.981 (0.947-1.015) |
| China Rural | 0.66 (0.39-1.11) | 0.954 (0.903-1.007) | 0.65 (0.35-1.18) | 0.989 (0.971-1.007) |
| Pooled fixed<br>effect | 1.02 (0.84-1.25) | 0.987 (0.960-1.015) | 0.80 (0.66-0.97) | 0.986 (0.977-0.995) |
| Heterogeneity | 0.0% | 0.0% | 0.0% | 0.0% |

1. Accounting for the competing risk of dementia-free death 2. Adjusted for age, gender and education level 3. Sub-hazard ratio

### Skull circumference and incident dementia

There was no evidence for an association between skull circumference and incident dementia in any site, or after meta-analysis, whether comparing quarters with the largest and smallest skulls (ASHR 1.02, 95% CI, 0.84-1.25, I^2^=0%), or the linear effect per centimetre of skull circumference (ASHR 0.987, 95% CI 0.960-1.015, I^2^=0%) (Table 2). The interaction between skull circumference (per quarter) and gender (male versus female) was statistically significant (pooled ASHR 0.86, 95% CI 0.75-0.98, I^2^=0.0%), but in the reverse direction to that hypothesized with a greater protective effect of larger skulls in men than in women. There was a non-significant trend towards an interaction between skull circumference and education (per level) (pooled ASHR 0.986, 95% CI 0.971-1.001, I^2^=32.1), also in the opposite direction to that hypothesized, with a greater protective effect at higher educational levels.

### Changes in leg length and skull circumference measures

Correlations between baseline and follow-up skull circumferences (0.53) and leg lengths (0.43) were generally modest. Correlations were low in the rural China site (0.20 and 0.21 respectively), and high in the urban Peru site (0.83 and 0.70 respectively).

Inspection of the distribution of change scores revealed some biologically implausible differences at the extremes of the distributions. Even after exclusion of the outliers (top and bottom 5% of the change score distributions), there was a tendency for both skull circumferences (mean difference −0.20, 95% CI −0.23 to −0.17 cms) and leg lengths (mean difference −0.99, 95% CI −1.12 to −0.85 cms) to be smaller on the second measurement occasion. Apparent shortening of leg length was more marked among those with dementia at baseline, than in those with CIND, and those who were cognitively normal (Table 3). Changes in skull circumference did not differ between those with dementia at baseline and those who were cognitively normal (Table 3).

**Table 3.** Changes in measurements of leg length and skull circumference, from baseline to followup, by baseline cognitive status (data pooled across sites)

|  | Change in leg length from baseline to follow-up |  | Change in skull circumference from baseline to follow-up |  |
| --- | --- | --- | --- | --- |
|  | N | Change score (95% CI) | n | Change score (95% CI) |
| 1. Cognitively normal | 6555 | -0.85 (-0.99 to -0.71) | 6760 | -0.20 (-0.23 to -0.16) |
| 2. CIND<sup>1</sup> | 366 | -1.31 (-1.87 to -0.75) | 375 | -0.08 (-0.23 to +0.07) |
| 3. Dementia | 442 | -2.67 (-3.18 to -2.17) | 480 | -0.33 (-0.45 to -0.20) |
| All | 7383 | -0.99 (-1.12 to -0.85) | 7615 | -0.20 (-0.23 to -0.17) |
| Overall effect of baseline cognitive status | F=21.1, 2 df, p<0.001 |  | F=6.6, 2 df, p=0.03 |  |
| Scheffe test | p-value for group comparison |  | p-value for group comparison |  |
| 3 versus 1 | p<0.001 |  | p=0.13 |  |
| 3 versus 2 | p=0.004 |  | p=0.04 |  |
| 2 versus 1 | p=0.34 |  | p=0.30 |  |

1. Cognitive Impairment, No Dementia

### Post hoc sensitivity analyses

Given modest correlations between baseline and follow-up measures of skull circumference and leg length, we repeated the regressions reported in Table 2, limited to those individuals with baseline and follow-up anthropometric assessments, and excluding the 10% with the most discrepant baseline and follow-up measures. This restriction did not affect the main findings (skull circumference pooled AHR 1.05, 95% confidence intervals 0.80-1.36, I^2^=37.9%; leg length AHR 0.76, 95% CI 0.60-0.97, I^2^=0.0%). Given the association between baseline dementia status and subsequent change in measured leg length, we assessed the association of change in leg length (per centimetre) with incident dementia, controlling for age, gender and education. The pooled effect size (AHR 1.006, 95% CI 0.992-1.020, I^2^=19.3%) did not suggest any association. Neither was change in skull circumference associated with incidence of dementia (pooled AHR 1.016, 95% CI 0.961-1.074, I^2^=7.7%). Since anthropometric measures were not available at follow-up on those who were deceased, these analyses could not take account of the competing risk of dementia free death, and were simple Cox’s Proportional Hazard regressions.

Finally, other than baseline cognitive status, physical frailty seemed to be the most prominent independent determinant of change in leg length, (adjusted mean difference, controlling for age, gender, education and CIND, −1.19, 95% CI −1.89 to −0.70 cms). We therefore repeated the regression for the effect of baseline leg length on incident dementia (reported in Table 2) controlling for frailty in addition to age, education and gender, and there was only a very small attenuation of the effect (ASHR 0.82, 95% CI 0.67-0.99, I^2^ = 0%).

## Discussion

### Main Findings

This study provides additional support for the salience of early life development to late-life dementia risk. However, while the hypothesis that longer legs are associated with a reduced risk of incident dementia was supported, the hypothesis that larger skulls would also be associated with a reduced risk was not. The association between longer legs and reduced dementia risk was independent of age, gender, education and physical frailty, and there was no heterogeneity of effect across sites. There was no evidence that the effect of leg length on incident dementia was modified by gender. The effect of skull circumference on incident dementia was modified by gender and educational level, but in the contrary direction to previous findings, and our hypotheses.

### Strengths and weaknesses of the study

The associations were assessed longitudinally, in large population-based dementia-free cohorts, comprising rural and urban catchment area sites in the Caribbean, Latin America, and China. Fixed effect meta-analysis was appropriately applied to increase the precision of our estimates, given the negligible heterogeneity for all of the associations studied. There have been only two previous longitudinal studies of these associations [14,17], neither of which approached the power and precision conferred by our 10,000 participants and 40,000 person years of follow-up.

A unique aspect of our approach was the repeated anthropometric measures at baseline and follow-up. The moderate agreement for the repeated measures, particularly for skull circumference suggested random measurement error, with potential for underestimation of the true effect of these exposures upon dementia risk. In a post-hoc analysis, the inverse association between leg length and incident dementia was more pronounced when excluding those with more discrepant baseline and follow-up leg length measurements, suggesting regression dilution bias. Random measurement error might have been reduced by more appropriate measurement methods (for example knee height [17] or sitting minus total height [32]), and more training and quality control. We also found a consistent tendency for both leg and skull dimensions to be smaller on the second measurement occasion. For legs, but not skulls, this was more pronounced for those with dementia at baseline than those who were cognitively normal. Although mortality was high among those with dementia [22], and those who died did not have a second anthropometric assessment, selective mortality seems an unlikely explanation. Leg lengths, per se, are unlikely to shorten other than after hip fracture. However, our measurement approach, measuring standing leg length, may have led to apparent shortening (and, in some cases, lengthening) due to difficulty in keeping the knee joint fully extended, particularly for those with arthritis, sarcopaenia, general fatigue, and, possibly, cognitive impairment. Physical frailty (capturing several of these elements) was associated with leg length shortening. Given these findings, the possibility of reverse causation, or residual confounding cannot be positively excluded. However, the association between baseline leg length and incident dementia persisted after controlling for physical frailty, and there was no association between change in leg length (or skull circumference) and incident dementia.

### Contextualisation with other research

The relationships between birthweight, subsequent growth, and cognitive development are well established, with possible ‘sensitive periods’ in infancy, childhood and adolescence [33,34]. In a recent analysis of historical cohort data from the UK MRC National Survey of Health and Development, trunk and leg length were inversely associated with cognitive function at age 53 years. These associations attenuated after controlling for adverse early life circumstances, and were entirely accounted for controlling for cognition at age 15 years [32]. Hence early-life skeletal growth may be a marker for cognitive development, and associations of adult leg length with late-life cognition and dementia risk may be accounted for by tracking of cognitive ability across the life course, and the cognitive reserve hypothesis, respectively. This would not, however, adequately account for the discrepancy, in our analyses, between leg length and skull circumference as predictors of incident dementia. Although adult leg length and height is mainly influenced by early growth and its determinants [35], legs continue to grow for up to a decade after skull growth is complete. Skull circumference and leg length may therefore be markers of different phases, and, possibly, different aspects of the developmental process. Shorter limb length is associated with cardiovascular risk factors in later life, particularly those linked to insulin resistance [36]. This may be another mediating pathway, best explored in birth cohorts with mid-life assessment of cardiovascular risk profiles.

Leg length, in contrast to trunk height, and skull circumference are thought to be stable across the adult life course. We found few previous studies of changes in measured leg length and skull dimensions in older populations. In a report from our 10/66 DRG site in Chennai, India where follow-up was limited to those with dementia or cognitive impairment, reassessment of skull circumferences and leg lengths three years after baseline indicated that leg lengths did appear to shorten, but at a similar rate in those with and without dementia, while skull circumferences remained stable [37]. In the context of dementia, in our baseline 10/66 DRG surveys, we found no cross-sectional associations between dementia severity and either leg length or skull circumference [18], and longitudinal computed tomography studies of people with dementia have not suggested that their cranial dimensions decrease with progression of the disease [12].

## Conclusions

Our cohort study clarifies the association between markers of early life development; leg length (an inverse association) and skull circumference (no association) and the incidence of dementia in late life. It addresses previous unresolved concerns regarding direction of causality through repeated measures of leg and skull dimensions. Main effects and interactions are estimated with much more power and precision than had been previously possible. Findings from previous studies in high income countries, can now be generalised to diverse urban and rural settings in Latin America and China.

Consistent findings across settings and previous studies provide quite strong support for an association between adult leg length and dementia incidence in late-life, with early life nutrition most plausibly implicated as the antecedent cause. While attention has focused upon education as a modifiable risk factor for dementia, much less has been accorded to other developmental factors. Mean male adult height, after a long period of relative stability, has increased by 5-10cms since 1920 in high income countries [38]. This secular trend may, in part, explain recent trends towards a declining incidence of dementia in high income countries, which are not adequately accounted for by improvements in education and cardiovascular risk profile [39-41]. Our findings would predict a 7% reduction in the incidence of dementia with each five centimetre increment in height. Further clarification of these associations could inform predictive models of future incidence, prevalence and numbers affected, and invigorate global efforts to improve childhood nutrition, growth and development.

## Authors’ contributions

All of the authors (MJP, DA, MG, YH, IZJ-V, JJLR, AS, ALS, MED, MMG, ZL, JJL-G, MAP) worked collectively to develop the protocols and methods described in this paper. MJP led the research group. JLR (Cuba), DA (Dominican Republic), IZJ (Puerto Rico), MG (Peru), AS (Venezuela), ALS (Mexico), and YH (China) were principal investigators responsible for the field work in their respective countries. MJP conducted the analyses, advised by MED, and wrote the first draft. All other authors reviewed the manuscript, provided further contributions and suggestions. All authors read and approved the final manuscript.

## Acknowledgements

The 10/66 Dementia Research Group’s research has been funded by the Wellcome Trust Health Consequences of Population Change Programme (GR066133 - Prevalence phase in Cuba and Brazil; GR080002- Incidence phase in Peru, Mexico, Cuba, Dominican Republic, Venezuela and China), the World Health Organisation (India, Dominican Republic and China), the US Alzheimer’s Association (IIRG - 04 - 1286 - Peru, Mexico and Argentina), and FONDACIT (Venezuela). The analysis reported here was carried out with funding support from the European Research Council (ERC-2013- ADG 340755 LIFE2YEARS1066). The Rockefeller Foundation supported our dissemination meeting at their Bellagio Centre.

